# Functional connectome fingerprinting accuracy in youths and adults is similar when examined on the same day and 1.5 years apart

**DOI:** 10.1101/812719

**Authors:** Maria Jalbrzikowski, Fuchen Liu, William Foran, Lambertus Klei, Finnegan J. Calabro, Kathryn Roeder, Bernie Devlin, Beatriz Luna

## Abstract

Pioneering studies have shown that individual correlation measures from resting-state functional magnetic resonance imaging studies can identify another scan from that same individual. This method is known as “connectotyping” or functional connectome “fingerprinting”. We analyzed a unique dataset of 12-30 years old (N=140) individuals who had two distinct resting state scans on the same day and again 12-18 months later to assess the sensitivity and specificity of fingerprinting accuracy across different time scales (same day, ~1.5 years apart) and developmental periods (youths, adults). Sensitivity and specificity to identify one’s own scan was high (average AUC=0.94), although it was significantly higher in the same day (average AUC=0.97) than 1.5-year years later (average AUC=0.91). Accuracy in youths (average AUC=0.93) was not significantly different from adults (average AUC=0.96). Multiple statistical methods revealed select connections from the Frontoparietal, Default, and Dorsal Attention networks that enhanced the ability to identify an individual. Identification of these features generalized across datasets and improved fingerprinting accuracy in a longitudinal replication data set (N=208). These results provide a framework for understanding the sensitivity and specificity of fingerprinting accuracy in adolescents and adults at multiple time scales. Importantly, distinct features of one’s “fingerprint” contribute to one’s uniqueness, suggesting that cognitive and default networks play a primary role in the individualization of one’s connectome.

## Introduction

Accumulating evidence from resting-state functional magnetic resonating imaging (rsfMRI) indicates that brain network architecture is highly individualized [Gordon et al., 2017; Gratton et al., 2018; Laumann et al., 2015a; Laumann et al., 2015b]. Improving our understanding of individual-level brain activity is leading to a mechanistic understanding of how neural activity contributes to individual differences, extending to individual differences in behavior. Furthermore, advances using individual-level neuroimaging markers to reflect indicators of pathology could, in the future, significantly improve our ability to make informed clinical decisions. A first step towards achieving this “personalized neuroscience” is to understand normative variation in individual-level features of brain activity, particularly through a developmental period when psychiatric illness typically emerges.

In participants with high-quality, densely sampled neuroimaging information, within-subject variability explained approximately one third of the variance in the data and is similar in magnitude to the explanatory power of the average or canonical network structure [Gratton et al., 2018; Power et al., 2011; Yeo et al., 2011]. Using alternative methods, others have also obtained similar estimates of intra- and inter-subject variation in rsfMRI data [Betzel et al., 2019; Mueller et al., 2013]. While these studies have produced important insights, such large amounts of neuroimaging data (~5-84 hours of data per individual, see [Gratton et al., 2019] for a review) may be infeasible for clinical samples.

In a related line of research, several studies show that individuals’ patterns from one scan can identify another scan from that same individual at a high level of accuracy [Finn et al., 2015; Horien et al., 2019; Miranda-Dominguez et al., 2014]. This method is known as “connectotyping” [Miranda-Dominguez et al., 2014] or functional connectome “fingerprinting” [Finn et al., 2015]. This analytic framework is effective with smaller amounts of rsfMRI data (3-15 minutes). Understanding the individual-level network measurement characteristics in a normative sample could provide a foundation for later parsing out meaningful differences in behavior and/or pathology.

Here, we aimed to build on initial findings indicating individualized brain functional architectonics [Finn et al., 2015; Miranda-Dominguez et al., 2014; Waller et al., 2017; Xu et al., 2016] to characterize how and which brain patterns are unique to an individual and to what extent this pattern is stable or changes over time. First, we assessed the sensitivity of fingerprinting measures, i.e. the likelihood of obtaining a true positive, and the specificity of these measures, i.e. the probability of distinguishing true negatives [Florkowski, 2008; Youngstrom, 2014]. To determine the specificity and sensitivity of fingerprinting, we applied a classification procedure. Scans from the same individual were viewed as positive pairs while all of the others were considered negative pairs. The area under the curve (AUC)-receiver operating characteristics (ROC) curve was utilized as a performance measurement to identify the sensitivity (true positive rate) and specificity (1-false positive rate) of fingerprinting at different thresholds and time scales. To measure group-level influences, we first identified predictive edges from a discovery sample and then examined to what extent these edges improved fingerprinting in a replication sample.

The majority of psychiatric disorders emerge during adolescence [Paus et al., 2008], a period of remarkable neuroplasticity and change [Larsen and Luna, 2018; Luna et al., 2015; Murty et al., 2016]. Thus, is it important to establish individualized brain markers of fingerprinting accuracy in this period and contrast them to that of adults. The extent to which there are age-associated differences in fingerprinting measures is an open question, given inconsistent findings in the literature [Demeter et al., 2019; Horien et al., 2019; Kaufmann et al., 2017]. One study used a combination of resting-state and task-based MRI scans to assess fingerprinting accuracy from same day scans and found that fingerprinting accuracy significantly improved from late childhood through adulthood [Kaufmann et al., 2017].. However, others found that fingerprinting accuracy is not affect by age in adolescents when scans were months to years apart [Horien et al., 2019] and fingerprinting accuracy in pediatric and adult samples is quite similar within a 5-18 month time frame [Demeter et al., 2019]. Insofar as we are aware, there has not yet been a direct comparison of scans completed on the same day versus those completed much later (i.e., a year), and to what extent this direct comparison changes or remains the same across adolescence. It is possible that the neuroplasticity observed in resting state scans during adolescence [Calabro et al., 2019; Jalbrzikowski et al., 2017] and/or motion artifact known to be more predominant in youth [Power et al., 2011; Satterthwaite et al., 2012]; reduces the ability to accurately identify an individual’s scan. Alternatively, “functional fingerprinting” of an individual’s resting state scan could be robust to these changes.

In our discovery sample, we leveraged a unique, two-time point data set that had two resting state scans from the same individual conducted on the same day (Visit 1, V1), and two resting state scans from the same individual collected on the same day 12-18 months later (Visit 2, V2; henceforth 1.5 years). We then assessed the level of sensitivity and specificity of fingerprinting accuracy; compared whether it is as stable for the same day as it is 1.5 years later; and determined if sensitivity and specificity at these different time scales were similar in youths and adults. We also used multiple statistical methods to determine connections that are “predictive” of individuals’ scans, reflecting one’s uniqueness. Finally, we explored how these edges performed in a replication sample.

## Participants

The final discovery sample consisted of 140 participants (1-2 visits, mean time between visits: 18 months, range of time between visits: 17-25 months). To test the generalizability of the predictive edges identified in our discovery sample, we tested the extent to which the previously identified features from each method improved fingerprinting accuracy in a replication sample with longitudinal data (N=208, 1-3 visits, mean time in between visits: 20 months, range of time between visits: 12 -50 months).

All participants were recruited from the greater Pittsburgh metro area. Participants and their first-degree relatives did not have a psychiatric disorder, as determined by phone screen and a clinical questionnaire. Exclusion criteria for all participants included any drug use within the last month, history of alcohol abuse, medical illness affecting the central nervous system function, IQ lower than 80, a first-degree relative with a major psychiatric disorder, or any MRI contraindications. There are previous publications using rsfMRI data from individuals within the replication sample addressing separate questions [Jalbrzikowski et al., 2019, in press; Jalbrzikowski et al., 2019; Marek et al., 2015].

Demographic information for both samples is reported in Table 1. Detailed inclusion/exclusion criteria are reported in the Supplemental Material.

**Table 1.** Participant Information for discovery and replication samples. Detailed exclusion criteria are provided in Figure S1.

| Discovery Sample |  |  |  |  |  |  |  |  |  |  |
| --- | --- | --- | --- | --- | --- | --- | --- | --- | --- | --- |
|  | Pre-task |  |  |  |  | Post-task |  |  |  |  |
|  | N | M/F | Mean Age [SD] | Age Range | Mean FD [SD] | N | M/F | Mean Age [SD] | Age Range | Mean FD [SD] |
| Visit 1 | 140 | 67/73 | 19.7 [5.0] | 12.0-31.0 | 0.24 [0.1] | 126 | 57/69 | 20.1 [4.9] | 12.0-31.0 | 0.25 [0.1] |
| Visit 2 | 93 | 48/45 | 22.2 [5.0] | 13.5-32.6 | 0.22 [0.1] | 86 | 45/41 | 22.3 [4.9] | 13.5-32.6 | 0.23 [0.1] |
| Replication Sample |  |  |  |  |  |  |  |  |  |  |
|  | N | M/F | Mean age |  | Age Range | Mean FD |  |  |  |  |
| Visit 1 | 208 | 104/104 | 18.5 [4.4] |  | 10.1-25.9 | 0.15 [0.05] |  |  |  |  |
| Visit 2 | 85 | 45/40 | 17.0 [2.8] |  | 11.9-21.8 | 0.12 [0.03] |  |  |  |  |
| Visit 3 | 59 | 32/27 | 18.2 [2.7] |  | 13.3-23.0 | 0.14 [0.03] |  |  |  |  |

## MR Data Acquisition: Discovery Sample

Data were acquired using a Siemens 3 Tesla mMR Biograph with a 12-channel head coil. Subjects’ heads were immobilized using pillows placed inside the head coil, and subjects were fitted with earbuds for auditory feedback to minimize scanner noise. For each rsfMRI run, we collected eight minutes of resting-state data, eyes open. Resting state data were collected using an echo-planar sequence sensitive to BOLD contrast (T_2_*). rsfMRI parameters were Repetition Time/Echo Time=1500/30.0 ms; flip angle=50°; voxel size = 2.3×2.3×2.3 mm. Structural images were acquired using a T1 weighted magnetization-prepared rapid gradient-echo (MPRAGE) sequence (TR/TE = 2300/2.98 ms; flip angle=9°; voxel size = 1.0×1.0×1.0 mm).

Participants completed a unique two-visit scan protocol. In the first visit, individuals participated in a MRI protocol (Visit 1, V1) that included two rsfMRI runs (Pre-Task, Post-Task), with an fMRI reward learning task (~40 minutes) conducted between these two runs. Approximately 1.5 years later, the same individuals returned and completed an identical MRI protocol (Visit 2, V2), which also included two rsfMRI runs (Pre-Task, Post-Task) separated by the same fMRI task. A visual depiction of the scan protocol, along with the respective names given to each run or scan, are presented in Figure 1.

**Figure 1.**
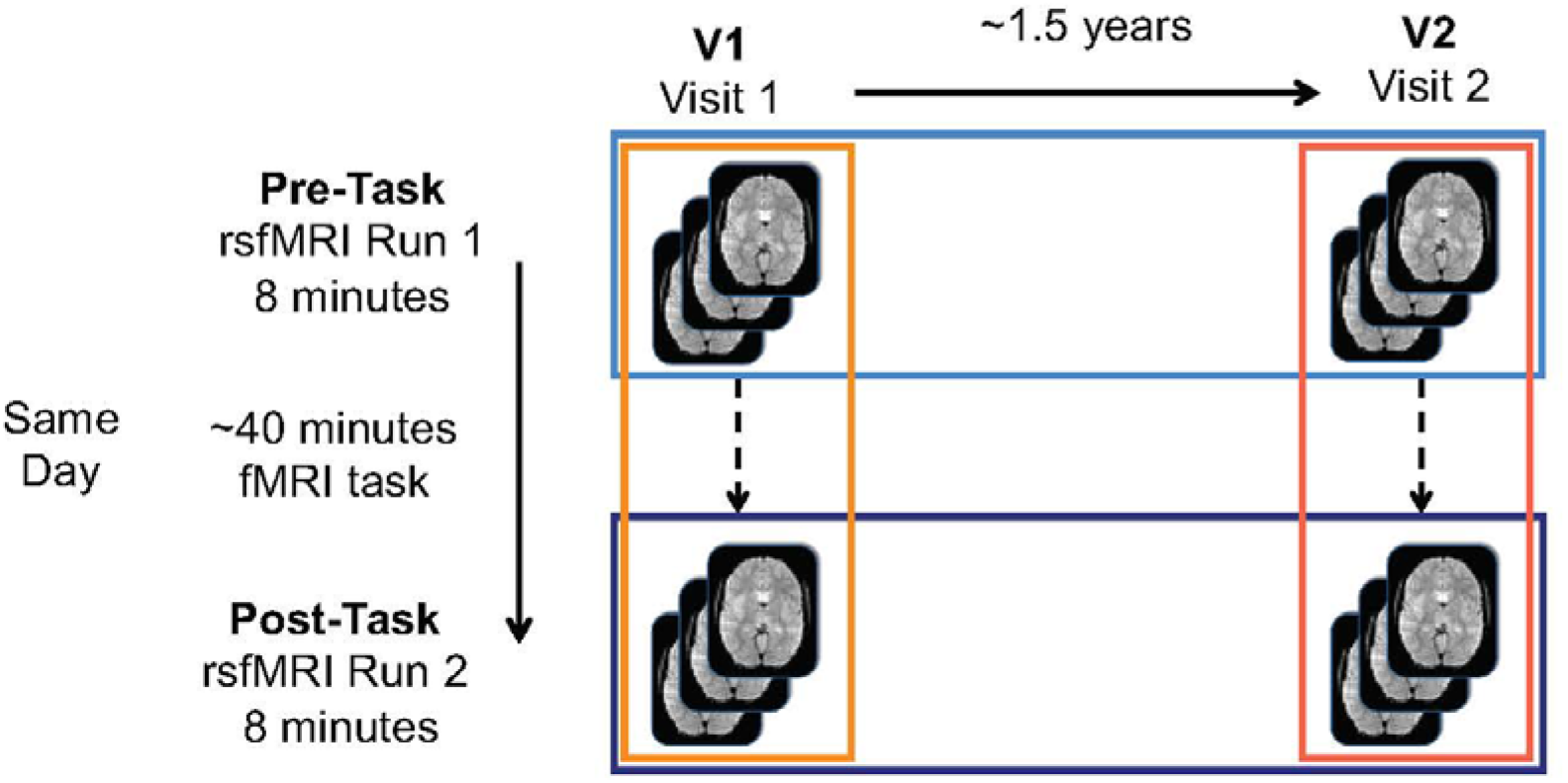
Visualization of protocol set-up. The colors refer to comparisons that will be made throughout the manuscript (oranges: same day comparisons, blues: 1.5 year comparisons).

## MR Data Acquisition: Replication Sample

Scan parameters for the replication sample are detailed in Supplementary text.

## rsfMRI Processing

First, we performed simultaneous slice-timing and motion correction of all functional images. Then, we implemented wavelet despiking to remove non-stationary events in the fMRI time series [Patel and Bullmore, 2015]. Third, functional images were warped into MNI standard space using a series of affine and nonlinear transforms. Then, after normalization, functional images were spatially smoothed using a 5-mm full width at half maximum Gaussian kernel. ICA-Aroma was then implemented to remove any remaining motion artifacts [Pruim et al., 2015a; Pruim et al., 2015b]. Finally, to control nuisance-related variability [Hallquist et al., 2013], we then conducted simultaneous multiple regression of nuisance variables and bandpass filtering at 0.009 Hz < *f* < 0.08. Nuisance regressors included were non-brain tissue (NBT), average white matter signal, average ventricular signal, six head realignment parameters obtained by rigid body head motion correction, and the derivatives of these measures. NBT, average white matter, and average ventricular signal nuisance regressors were extracted using MNI template tissue probability masks (>95% white mater, >98% cerebral spinal fluid, [Fonov et al., 2009].

For all subjects, we calculated a quality control measure with respect to head motion, namely volume-to-volume frame displacement (FD). Consistent with recent developmental cognitive neuroscience publications [Bathelt et al., 2019; Calabro et al., 2019; Hafeman et al., 2019; Li et al., 2019], subjects were removed from rsfMRI analyses if the average frame displacement across the run was > 0.5mm (N=5).

## Functional Network Parcellation

We applied a previously-defined, functional connectome parcellation of 333 functional regions of interest (ROIs) across cortical structures [Gordon et al., 2016] to each participant’s rsfMRI data (Figure 2A). This parcellation consists of 13 reliable rsfMRI networks, many of which have been identified in other studies, including the Frontoparietal, Default, and Visual networks [Glasser et al., 2016; Power et al., 2011; Shen et al., 2013]. See Supplementary Table S1 for a list of 13 networks and details about them (for each network: number of nodes, number of within-connectivity edges, and number of between-connectivity edges).

**Figure 2.**
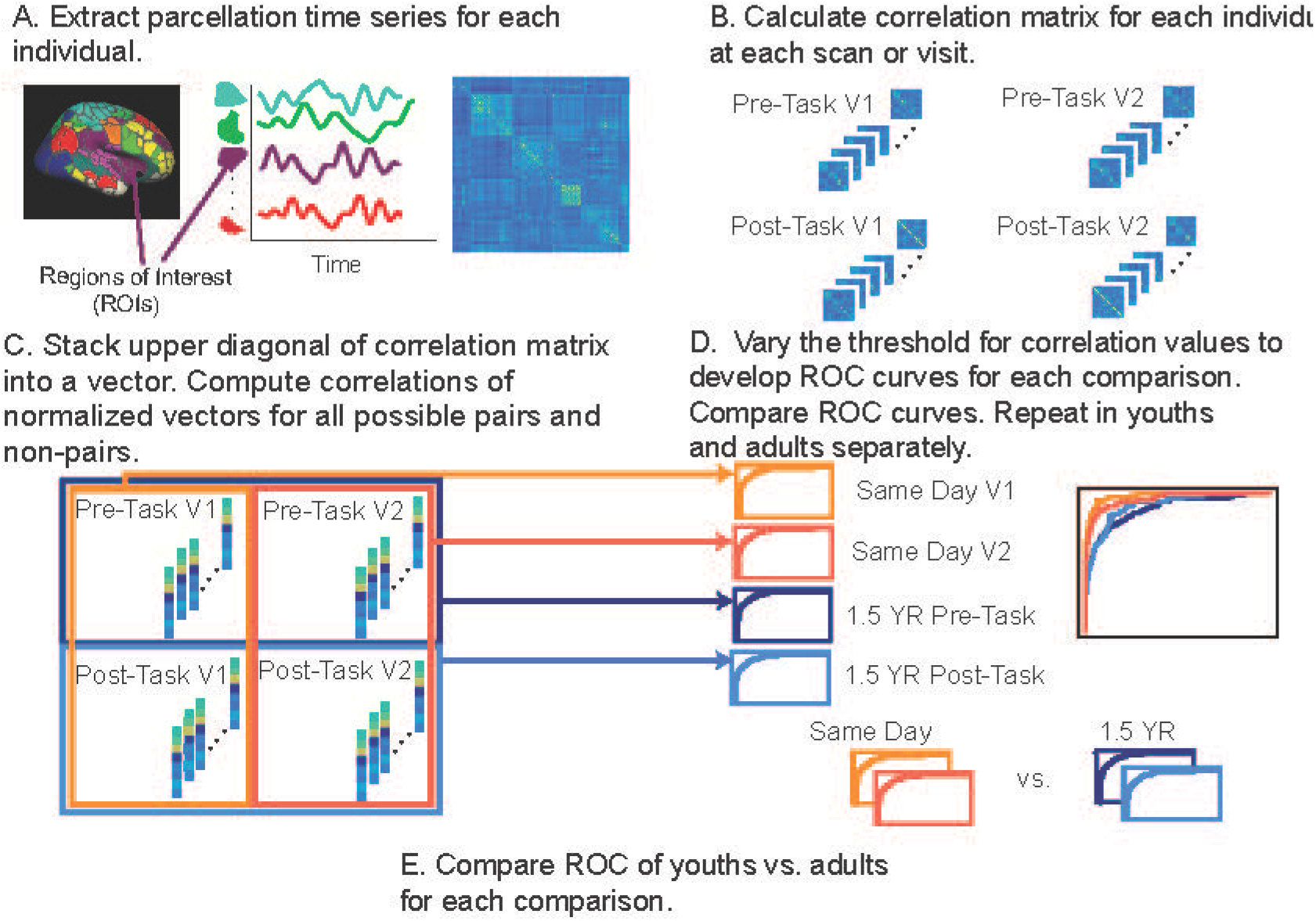
(A) After resting-state fMRI data were processed, we extracted out the time series from an established parcellation and (B) calculated a correlation matrix for each individual and their respective scan visit. (C) For each scan at each visit, we stacked a vector from the upper diagonal of the correlation matrix. Each stacked vector represents a scan from one person. The stacked vectors could be a separate scan from the same individual or a separate scan from a different individual. We computed correlations between each vector for all possible pairs and non-pairs. (D) By varying the threshold of the correlation values to determine what was a true-or false-positive, we developed ROC curves for each comparison, with a total of four comparisons: same day visit 1, same day visit 2, 1.5 years apart pre-task 1.5 years apart post-task. We then used DeLong’s method to compare the ROC curves. (E) For each of the four comparisons, we compared the ROC curves of youths vs. adults.

For each participant, we computed Pearson’s correlation of each ROI’s time series with that of every other ROI, producing a 333 x 333 correlation matrix (Figure 2B). The upper diagonal of the correlation matrix for each individual was stacked into a vector (55278 edges), and each vector was normalized to a mean of zero and variance of one (Figure 2C). We performed this procedure for each subject’s rsfMRI run, resulting in four normalized vectors for the majority of participants. The correlation between any two of these normalized vectors, *V* and *U*, was their dot product:

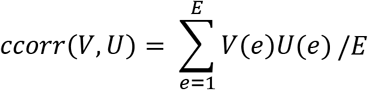

where *E* is the total number of edges and *e* is an individual edge.

### Identification Accuracy

Next, we sought a classifier to identify resting state fMRI-measured connectomes that “match”; ideally these would be connectomes from the same subject. To do so, we first computed the correlation of normalized vectors, *V* and *U*, for all possible pairs of subjects, as described above (Figure 2C). The classifier seeks a threshold *t* for these correlation values that yields a high rate of true positive identification, namely connectomes from the same subject, while minimizing the number of false positive identifications, connectomes from different subjects labeled as from the same subject. We varied *t* from interval of zero to one to create a receiver operating characteristic (ROC) curve. We then estimated the area under the curve to determine the accuracy of the classifier (Figure 2D). The *t* that maximized true positive rate-false positive rate (TPR-FPR) was chosen for reporting. ROC curves were generated for the entire sample for 1) same day identification accuracy (Pre-Task vs. Post-Task) and 2) identification accuracy 1.5 years apart (V1 vs. V2). To compare identification accuracy for same day vs. 1.5 years later, we compared ROC curves to each using DeLong’s test for two ROC curves ([DeLong et al., 1988; Robin et al., 2011], Figure 2D).

To determine whether fingerprinting accuracy was affected by age, we split the entire sample of 12-30 year olds by the median age (20.4 years), considering the participants “youths” if they were under the median age (V1 N=70) or “adults” if they were over the median age (V1 N=70). We then calculated ROC curves for youths and adults separately for 1) same day fingerprinting (Pre-Task vs. Post-Task) and 2) fingerprinting 1.5 years apart (V1 vs. V2). To test for significant differences in identification between youths and adults, we used DeLong’s test for two ROC curves (Figure 2E). We also assessed the effects of sex on fingerprinting accuracy by calculating ROC curves for each sex separately for each visit and session.

### Identification of Predictive Edges

Next, we sought to determine which edges contribute most to fingerprinting accuracy. We made four comparisons to assess identifiability of each subject (V1, pre versus post; V2, pre versus post; V1 pre versus V2 pre and V1 post versus V2 post) and the same comparisons among non-pairs, resulting in 98790 comparisons (same subject pairs=532, non-pairs=98258). Because there was an over-representation of non-pairs (two scans, one from individual *s* and another from individual *j*), we used synthetic minority over-sampling technique to reduce bias when selecting the predictive features [Chawla et al., 2002] and selected 532 pairs and 1596 non-pairs. This method uses Euclidean distance to select non-pairs closest to the pairs (Chawla et al., 2002). We then split the discovery sample into a training (2/3 of the data: 355 pairs, 1064 non-pairs) and test set (1/3 of the data: 177 pairs, 532 non-pairs). For each method, described below, we used 10-fold cross validation to identify the number of edges that gives the highest AUC. The terms of the dot product (i.e., from 52,278 edges) for each comparison were the input features.

#### Finn Method

We used the method that Finn et al. described a method to calculate the most predictive edges [Finn et al., 2015]. We then ranked all edges from most predictive to least predictive and we successively decreased the threshold of “most predictive” edges to develop the ROC curve. Through cross-validation in the training set, we determined that the optimal TPR-FPR rate was when we included the five percent “most predictive” edges. Then, in the test data set, we selected only these predictive edges and calculated a correlation between each scan and all other scans (sum of the dot product). We then used the correlation threshold to develop the ROC curve in the test data set.

#### Support Vector-Machine Learning and Elastic Net Regression Methods

Both of these methods are common tools for model selection. Here, the goal was to choose a set of predictive edges that differentiated same-subject pairs from other pairs. For both methods, the input information was the terms of the dot product between two scans. We developed optimal tuning parameters in the training data set and obtained weights for the selected edges. Elastic net regression was implemented using R package glmnet [Friedman et al., 2009]. The support-vector machine (SVM) analysis was implemented in R package, sparseSVM ([Yi and Zeng, 2018], with an elastic net penalty. For both methods, we implemented fivefold cross-validation and the tuning parameters were chosen from this procedure. We set *α* = 0.1 and chose the penalty parameter (*λ*) with the minimal cross-validation error. Within the discovery sample, we then applied the model developed in the training set to the test data set. By weighting the individual product of the selected edges, we obtained a predicted value, 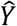, for which 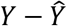 ranges from −1 to 1. We developed ROC curves by changing the threshold of 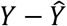 to determine what constituted a “pair” in the test data.

### Assessing over-representation of network connectivity in predictive edges

To identify network representation in the predictive edges for each method identified above, we conducted Chi-square tests to assess whether there was over-representation of 1) between (e.g. Frontoparietal-Default edges; the off-diagonal edges) or within-network connectivity (e.g., Frontoparietal-Frontoparietal edges; the block structure on the diagonal) and 2) specific within-network connectivity networks, and/or 3) distinct between-network connections. We used the standardized residuals (>3.0) to determine networks that contributed to one’s uniqueness.

### Performance of predictive edges in replication sample

To test the generalizability of the predictive edges identified in our discovery sample, we tested the extent to which the previously identified features from each method improved fingerprinting accuracy in a replication sample with longitudinal data.

### Effects of Possible Confounds

Because motion is a well-known confound in rsfMRI studies of development [Satterthwaite et al., 2012], we calculated multiple measures of motion: framewise displacement (FD) as described by Power et al [Power et al., 2012], as well as mean head displacement, maximum head displacement, the number of micro movements (N > 0.1 mm), and head rotation as described by Van Dijik et al [Van Dijk et al., 2012] we assessed whether motion differs between sessions or visits by running within-subjects t-tests of the motion variable between session (pre-, post-task) and visit (V1, V2). To assess the effects of age on motion, we conducted an independent samples t-test between group (youths 12.0-20.39 years old vs. adults 20.4-30 years old) for each session and visit.

Because “same day” scans were acquired within the same MRI session, improved same day fingerprinting accuracy could be due to better-quality registration from the same scan session. A subset of the participants (ages 18-30 years, N=76) participated in an additional scan on the same day, but in a different scan session (i.e, the participant came out of the MRI scanner and a few hours later participated in another MRI session). This subset of participants participated in a position emission tomography study that included an additional MRI scan with each visit. Classification as detailed previously was performed, using the separate scans at each visit to predict identification accuracy from this additional MRI session. See Supplementary Table S2 for details on this subset of participants.

## Results

Participant information for both visits is presented in Table 1. Participant information for youth and adult groups are reported in Supplementary Table S3.

### Same Day vs. 1.5 years later

Across the entire sample, identifiability on the same day was quite high (Average AUC: 0.94). Identification accuracy between scans 1.5 years apart was also high (Average AUC: 0.91, Figure 3A). However, same day accuracy was significantly higher than identification of scans 1.5 years apart (Figure 3A, Table 2A). Differences in fingerprinting accuracy on the same day compared to scans 1.5 years apart were observed for both youth (Figure 3B, Table 2B) and adult groups (Figure 3C, Table 2C).

**Table 2.**
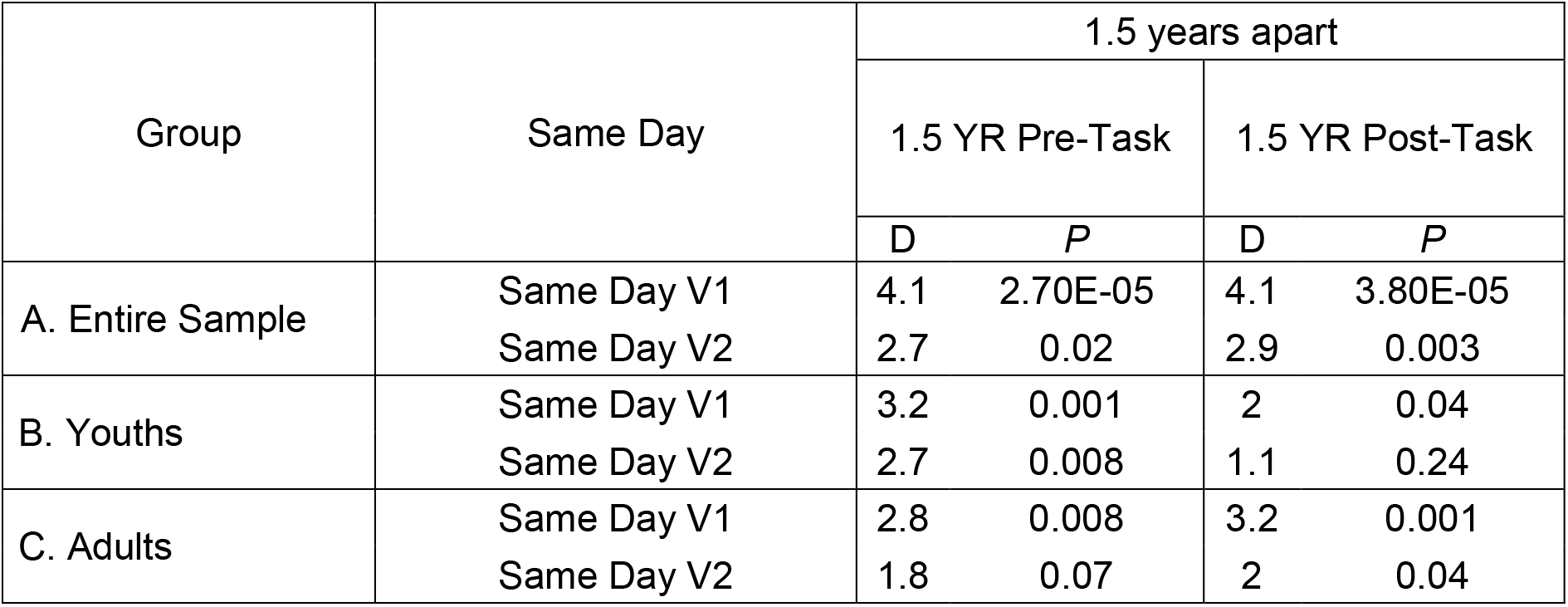
A) Across the entire sample, same-day identification accuracy was significantly higher than identification accuracy 1.5 years apart. The D-value represents the test statistic for the respective comparison. This pattern remained when youth (B) and adults (C) were assessed separately.

| Group | Same Day | 1.5 years apart |  |  |  |
| --- | --- | --- | --- | --- | --- |
|  |  | 1.5 YR Pre-Task |  | 1.5 YR Post-Task |  |
|  |  | D | P | D | P |
| A. Entire Sample | Same Day V1 | 4.1 | 2.70E-05 | 4.1 | 3.80E-05 |
|  | Same Day V2 | 2.7 | 0.02 | 2.9 | 0.003 |
| B. Youths | Same Day V1 | 3.2 | 0.001 | 2 | 0.04 |
|  | Same Day V2 | 2.7 | 0.008 | 1.1 | 0.24 |
| C. Adults | Same Day V1 | 2.8 | 0.008 | 3.2 | 0.001 |
|  | Same Day V2 | 1.8 | 0.07 | 2 | 0.04 |

**Figure 3.**
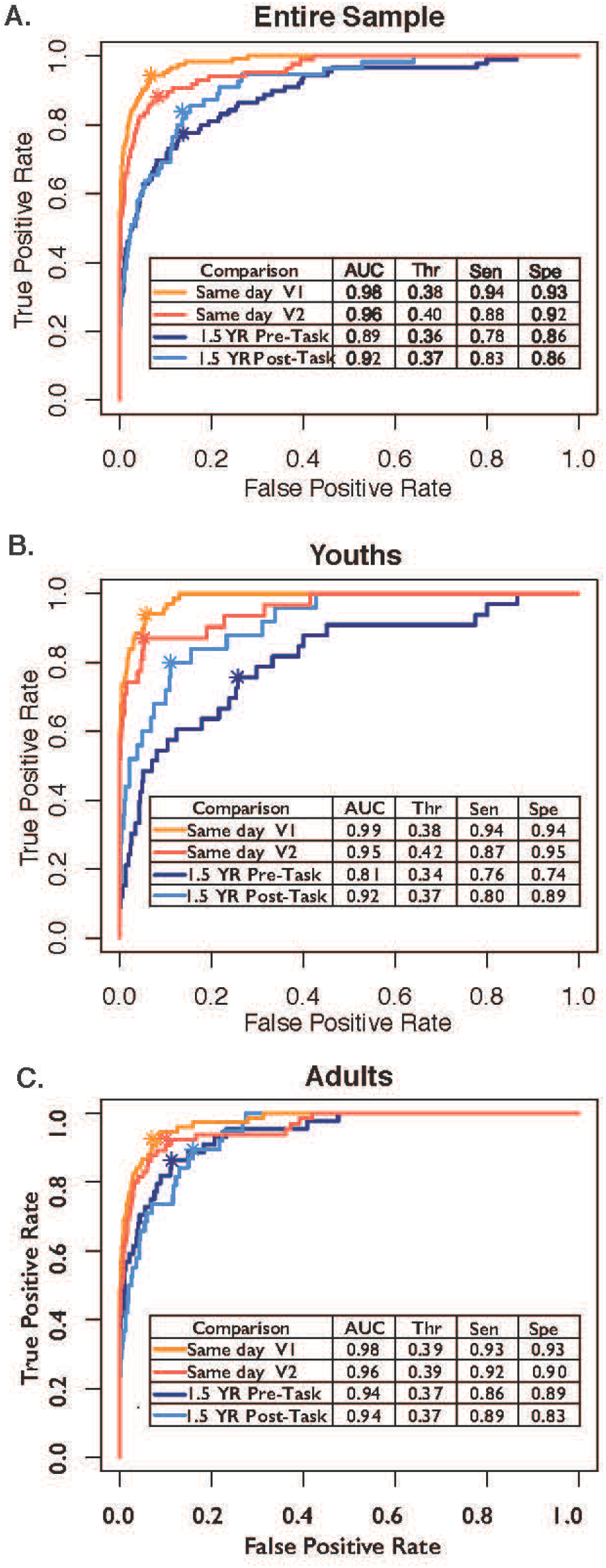
A) Across the entire sample, identification accuracy was higher for same day (orange) vs. 1.5 years later (blue). This pattern of results remained when youth (B) and adults (C) were assessed separately. The asterisk in each for each curve refers to the point when of optimal trade-off between sensitivity and specificity. AUC= Area under the curve, Thr=threshold at which optimal true positive rate was obtained. Sen=sensitivity, Spec=specificity

### Fingerprinting accuracy of youth vs. adults

Same day accuracy was not statistically different between youths and adults. The two groups exhibited similar levels of same-day identification accuracy (Table 3A). The same pattern of results emerged when comparing youths and adults on fingerprinting accuracy 1.5 years apart for three out of four of the comparisons (Table 3B). ROC Curves are presented in Figure 3. We also found a similar pattern of results when we split youths and adults by the mean age (20.8 years, 5 participants changed age group membership) and when we considered individuals as “adults” at age 18 (21 participants changed age group membership). Finally, we obtained a comparable pattern of results when we tested fingerprinting accuracy in males and females separately (Supplementary Table S3).

**Table 3.** A) Same-day fingerprinting accuracy is similar for youth and adults. B) Fingerprinting accuracy 1.5 years apart is similar for youth and adults in three out of four comparisons.

| Comparison |  | Adults vs. Youth |  |
| --- | --- | --- | --- |
|  |  | D | P |
| A. | Same Day V1 | -1.7 | 0.08 |
|  | Same Day V2 | 0.16 | 0.87 |
| B. | 1.5 YR: Pre-task V1 vs. Pre-task V2 | -2.7 | 0.006 |
|  | 1.5 YR: Pre-task V1 vs. Post Task V2 | -0.42 | 0.67 |
|  | 1.5 YR: Post-task V1 vs. Post-task V2 | -0.7 | 0.46 |
|  | 1.5 YR: Post-task V1 vs. Pre-Task V2 | -1.7 | 0.09 |

### All model selection methods tested improve identification accuracy in the test portion of the discovery sample

As seen in Figure 4A, the fingerprinting accuracy significantly improved when we used predictive edges selected by the Finn method, SVM, or Elastic Net to predict identification accuracy. The three model selection techniques performed similarly to one another (Supplementary Table S4). When we used only the predictive edges identified by these methods to re-assess fingerprinting accuracy in all previous comparisons, each method improved fingerprinting accuracy but did not change reported results.

**Figure 4.**
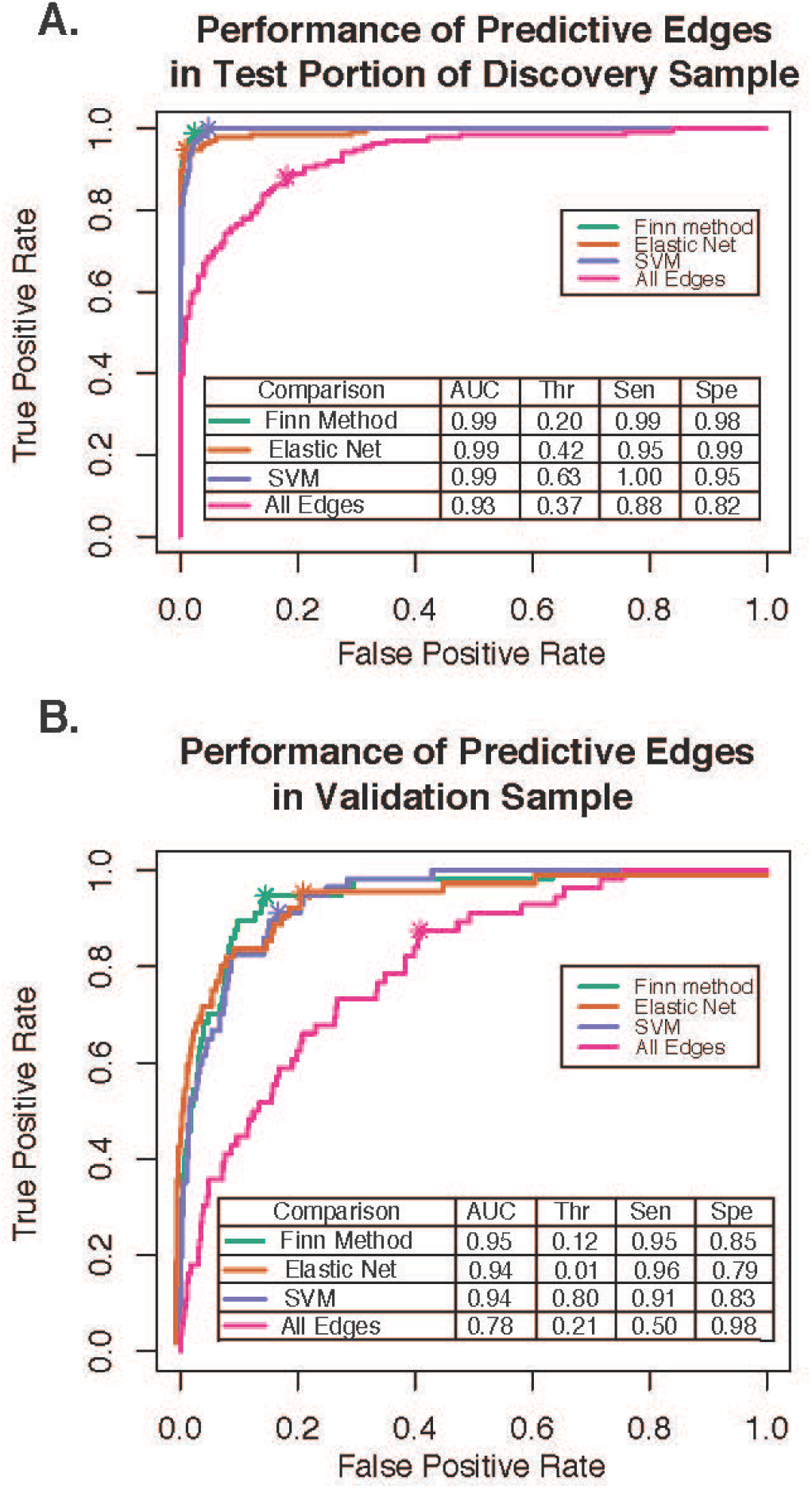
A). When edge selection was performed via various methods (Finn method (green), Elastic Net (orange), and SVM (purple)), identification accuracy was significantly improved in comparison to using all edges (pink) for identification accuracy. B. When we applied predictive edges previously identified in the training sample to an independent sample, all methods significantly improved identification accuracy.

### Predictive edges from discovery sample improves accuracy in replication sample

Similar to previous results [Finn et al., 2015; Waller et al., 2017], we found lower identification accuracy in a replication sample with more “standard” MRI parameters (e.g., longer TR, shorter length of scan, fewer number of head coils, Figure 4B, pink ROC curve). When we applied the most predictive edges from previously applied model selection techniques, however, fingerprinting accuracy significantly improved (Figure 4B, Supplementary Table S5). When we omitted pairs from scans conducted on the same day in the discovery sample and focused our analyses on identifying predictive edges from pairs that were 1.5 years apart, we obtained the same pattern of results (Supplementary Table S6). The weights for predictive edges from Elastic Net and SVM are provided in Supplementary Tables S7-8.

Optimal thresholds were determined separately in the discovery and validation sample prior to generating an AUC. This is necessary because of differences in the nature of the data. As shown in Figure S2, the distributions of the correlations between pairs were quite different in the two samples, precluding use of the selected discovery threshold for the validation threshold. Despite these differences, when the predictive edges derived from the discovery sample were applied to the validation sample, fingerprinting accuracy improved, suggesting that edges important for prediction are similar across samples.

### Predictive edges are over-represented in Frontoparietal, Default, and Dorsal Attention Networks

Figure 5A-C shows the relative contribution, normalized for number of edges in each network (or off/diagonal edge group) and number of edges determined to be predictive by each method. Across all methods, in comparison to between-connectivity edges (e.g., Frontoparietal-Visual connections, Supplementary Table S9), within-connectivity edges (e.g., Frontoparietal-Frontoparietal connections) were relatively more important in predicting identification accuracy. More specifically, within-connectivity edges from the Frontoparietal, Default, and Dorsal Attention networks drove this finding (Figure 5B). These networks had standardized residuals greater than 3.0 in all comparisons (Supplementary Table S10). A With SVM and Elastic Net, edges from the Ventral Attention network were also considered important predictors for fingerprinting accuracy. similar pattern emerged when we examined same-day and 1.5-year comparisons separately (Supplementary Table S10) and youth and adults separately (Supplementary Tables S12-13).

**Figure 5.**
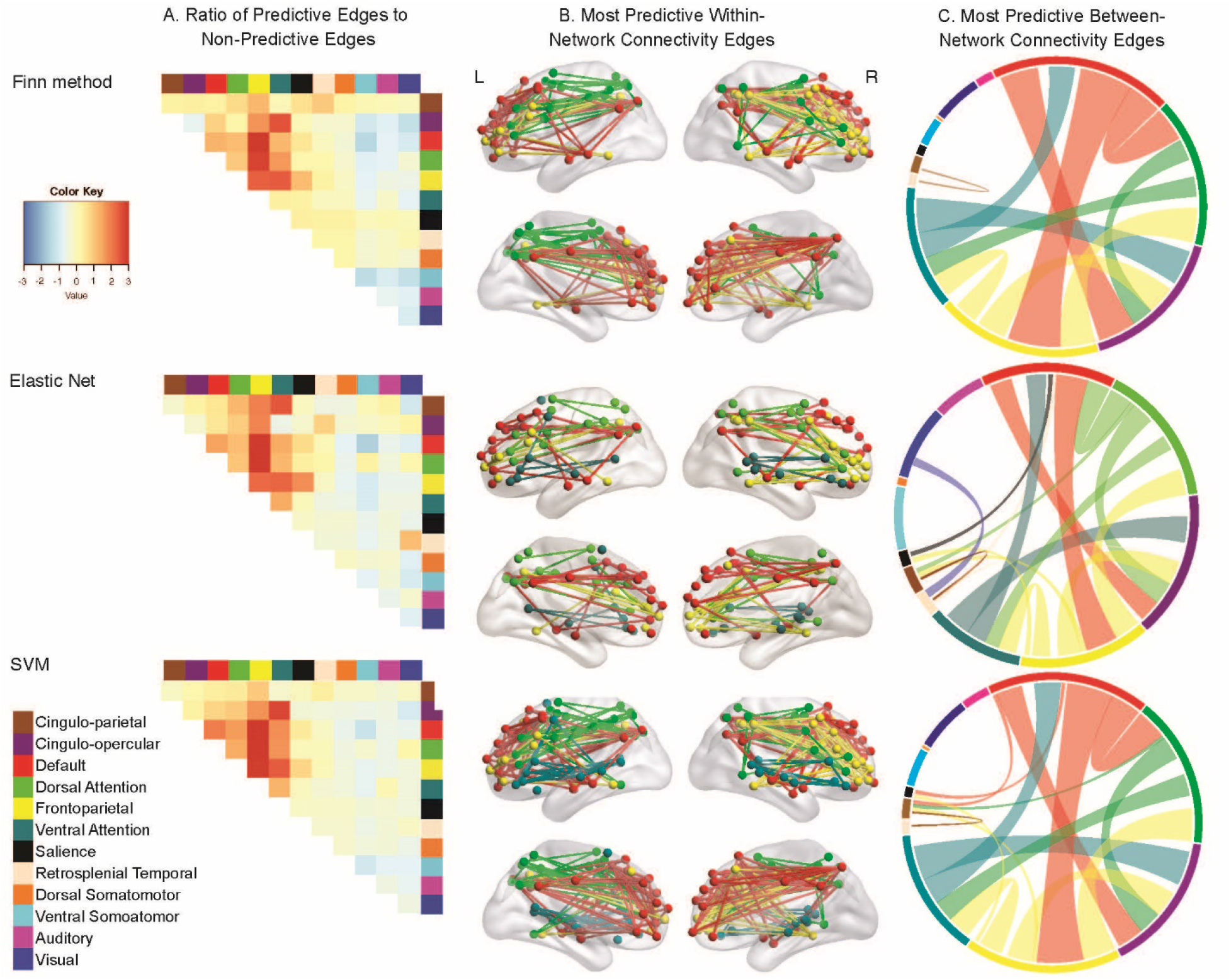
A) The ratio of predictive edges to non-predictive edges in each network connection, normalized for total number of edges in each connection for the three different methods. Warmer colors on the heatmap indicate that edges from a particular network are more important for identification accuracy. B) Within-network connections that are particularly important for identification accuracy in all three methods examined. In all methods, within connectivity edges in Frontoparietal (yellow), Default (red), and Dorsal Attention (bright green) networks are considered predictive. In the Finn method and SVM, within connectivity connections in the Ventral Attention network were also predictive of identification accuracy. C) Between-network connections that are particularly important for identification accuracy. The colors around the circle reflect the different networks examined. Thicker bands of color indicate a greater number of edges from that particular network were considered predictive. The between connectivity edges (lines going across the circle) were randomly chosen from one of the two connected networks (e.g., Between network connectivity between the Default and Cingulo-Opercular network is red, but between network connectivity between the Default and Ventral Attention network is green).

We also examined which between-network connectivity edges were relatively important for fingerprinting accuracy. Across all three methods, connections between the Fronto parietal-Default, Frontoparietal-Dorsal Attention, Ventral Attention-Cingulo-opercular, and Frontoparietal-Ventral Attention networks were over-represented in comparison to other connections (Figure 5C). See Supplementary Table S14 for standardized residuals from chi-square test of between-network connectivity and Supplementary Table S15 for standard residuals from chi-square test of over-representation of all network connections. Similar patterns emerged when we examined same-day and 1.5-year comparisons separately (Supplementary Table S16) and youth and adults separately (Supplementary Table S17).

### Testing Effects of Possible Confounds

To ensure that the improved same-day accuracy was not because individuals were in the same scan session and benefiting from improved MRI registration, we examined a subset of individuals (N=76) who completed an additional MRI session on V1 and V2 after being taken out of the scanner and repositioned. Supplementary Table S18 reports similar AUC, threshold levels, and sensitivity/specificity for identification accuracy within the same MRI session (V1 Pre-Task predicting accuracy of V1-Post-Task) and different MRI session on the same day (e.g., V1 Pre-Task predicting identification of extra session V1).

When we compared head motion between sessions, one out of two of the session comparisons (pre-task V1 versus post-task V1) was statistically significant. Individuals had higher FD during post-task V1 in comparison to pre-task V1; *t*=-2.5, *p*=0.01). One out of two of the visit comparisons (post-task V1 vs. post-task V2) was also statistically significant. FD at post-task V2 was lower in comparison to FD at post-task V1 (*t*=2.8, *p*=0.006). Notable, while FD was elevated in post-task V1 in comparison to other sessions/visits, the impact of visit and session on fingerprint accuracy was similar in other comparisons, suggesting that motion differences were not driving our results. A similar pattern of findings was observed in other motion metrics (Supplementary Table S19).

There were no statistically significant differences between youths and adults at any session or visit in FD, mean head displacement, maximum head displacement, or number of micromovements (all p≥0.05). Results are reported in Supplementary Table S19.

To ensure that our results were not driven by parcellation choice, we also re-ran all analyses after extracting ROIs from two separate parcellations (Power et al., 2011; Shen et al., 2013) and obtained a similar pattern of results.

## Discussion

For neuroimaging to have clinical utility, it is essential to understand what to expect from an individual’s network characteristics in multiple contexts. Here we show that identification accuracy of one’s resting state scan – how much it reflects a “functional fingerprint” – depends on the timespan between assessments. We provide supporting evidence that adolescents have similar levels of fingerprinting accuracy to adults when visits are years apart [Horien et al., 2019; Miranda-Dominguez et al., 2018] and extend this literature to show that this finding is consistent on a same-day visit. Furthermore, we used multiple methods to identify a small number of edges consistently predictive of an individual’s scan. These edges are more likely to be in the Frontoparietal, Default, and Dorsal Attention networks. We identified these edges in a discovery sample and then used these edges to improve identification accuracy in a replication sample. We propose that particular edges in the Frontoparietal, Default, and Dorsal Attention networks contribute to an individual’s “uniqueness” and are similar in youths and adults. These results bring us a fuller understanding of functional networks in the human brain.

### Stability of identification accuracy across time

Our results indicate a high level of subject identification accuracy even after 18 months. although the greater time interval incurred a significant decrease in identification. This result provides compelling evidence that there are extant foundational properties to individualized resting state network organization that are persistent and specific to each individual. The significant degradation in identification after 18 months is not driven by registration, which we tested and found to not be a contributor. Reduced fingerprinting accuracy with time could reflect greater noise between two scans; alternatively, or in addition to, the small but significant degradation in identification could reflect inherent plasticity in network organization in both youths and adults.

### Networks that Underlie Prediction

We found evidence that identification was driven by edges particularly in the Frontoparietal, Dorsal Attention, and Default mode networks and were consistently identified by different analytical approaches. This is a striking result that identifies networks critical for higher-order cognitive processing and endogenous self-referential processing. Thus, these results provide suggestive evidence that how we engage foundational cognitive and endogenous processes may contribute to demarcating individuality. This finding also has the potential to inform our understanding of impaired development in psychopathology that may be particularly represented in these networks.

### Predictive edges from the discovery sample improve fingerprinting accuracy in the validation sample

We also show that predictive edges identified in one sample can be applied to an independent sample to improve identification accuracy, even then the validation sample has somewhat different properties than the discovery sample (Figure S2). Sets of edges making dominant contributions to fingerprinting accuracy vary amongst individuals. Though the original goal of fingerprinting accuracy was, in part, to identify the specific predictive edges to identify specific individuals [Finn et al., 2015], we implemented a different approach. In dermal fingerprinting, approximately thirty islets or forks on the ridges or “identification points” are used to demonstrate uniqueness [Galton, 1892; Stigler, 1995]. It is highly unlikely that any other individual will have the same patterns from combination of these distinct ridges. By showing predictive edges from one data set improve fingerprinting accuracy in an independent data set, we demonstrate that a similar phenomenon is taking place with rsfMRI fingerprinting.

Multiple methods performed quite well in the discovery sample; however, as is consistent with the literature, a reduction in sensitivity and specificity in the replication sample suggests some over-fitting in this sample. On the other hand, our replication sample is a strength of our study, because the majority of fingerprinting accuracy studies have not tested the utility of predictive edges in another data set. The robustness of these results demonstrates that particular edges carry the most information for identifiability. This is a first, important step for understanding how fingerprinting accuracy can be used in a clinical context. However, as observed in the two separate samples, the distributions of correlations for pair identification between the two samples are remarkably different (Figure S2). In line with other fields of medicine that use a biological measure to detect risk or disease status [Greenwood et al., 2012], it will be important to determine the quality of the rsfMRI scans to ensure valid analyses.

The lower sensitivity and specificity in our replication sample is consistent with reports of fingerprinting accuracy in lower quality rsfMRI protocols [Finn et al., 2015; Waller et al., 2017]. A number of factors could be contributing to the degradation in fingerprinting accuracy, including a shorter scan time [Finn et al., 2015] or decreased resolution of the rsfMRI scan. Additionally, the discovery sample had higher resolution structural scans to assist with accurate registration of the rsfMRI data. Partial volume effects could also be contributing the degradation in fingerprinting accuracy. In the future, it will be informative to test if incorporating methods to correct for partial volume effects [Dukart and Bertolino, 2014] will improve rsfMRI fingerprinting accuracy in less optimized rsfMRI protocols.

### Identification accuracy is similar in youths and adults

We did not observe differences in the fingerprinting accuracy between youth and adults, for both same day and 1.5-year comparisons. Indeed, others have recently found that identification accuracy is similar in youth and adults [Demeter et al., 2019; Horien et al., 2019]. We extend these findings by showing accuracy is high both for same day and longer-term (1.5 years) intervals in the same sample. Furthermore, because identification accuracy in both youth and adults was lower across a longer period of time and similar edges contributed to same day and 1.5-year identification accuracy, our results suggest that this reduction is not solely due to known developmental changes. Significant cognitive development occurs through adolescence [Larsen and Luna, 2018; Luna et al., 2015; Steinberg, 2005], in the context of evidence for stability at the group level in network properties [Hwang et al., 2013; Jalbrzikowski et al., 2019; Marek et al., 2015; Marek et al., 2019]. The stability in identification accuracy across development further supports that implication that network properties contain individualized foundational properties that define uniqueness.

In one case, we did find that youths had significantly worse identification accuracy in comparison to adults (Table 3B: 1.5 YR: Pre-task V1 vs. Pre-task V2) when the scans were 1.5 year apart. However, in three out of four similar relevant comparisons, we did not observe differences in fingerprinting accuracy between youths and adults. In this particular comparison (i.e., Pre-Task V1 to Pre-Task V2), reduced accuracy in these youths was driven by the lower identification accuracy in the pre-task scans: we speculate that youths are more variable and excitable when first getting in a scanner, perhaps because they have had less experience with life events akin to an MRI scan than do adults. Furthermore, we know that identification accuracy is reduced with increased head motion [Horien et al., 2018], and youth have greater levels of head motion in comparison to adults [Satterthwaite et al., 2012]. However, in our sample, we did not see a statistical difference between head motion in these two groups and our results remain stable when we use more conservative framewise displacement thresholds (FD < .3).

### An Improved Statistical Framework for Identification Accuracy

We show that edges important for identification accuracy are similar across different methods used to identify them. Furthermore, predictive edges identified in one sample can be applied to an independent sample to improve identification accuracy. The robustness of these results demonstrates that particular edges carry the most information for identifiability.

We also believe this statistical framework also improves upon the majority of previous methods used in this area. Previous methods, for instance, presume there is a “match” for each respective scan in the data set (i.e., each individual has at least two scans in the pool of available data) and do not consider false positives. Furthermore, given that research shows that identifiability of individuals decreases as the sample size increases [Waller et al., 2017], it is important to account for sample size in the model. Additionally, while many studies show that the identification test metric for individual identifiability is significantly greater than would be expected by chance, it is difficult to know how meaningful this metric is when identification accuracy is in the range of ~40-60% (e.g., [Horien et al., 2019]. Additionally, we were interested in understanding how feature selection approaches (e.g., elastic net and SVM) compare to the original method presented by Finn and colleagues (2015). Similar to our approach, SVM has also recently been successfully applied to better understand genetic similarity in fingerprinting accuracy [Demeter et al., 2019]. Finally, we assured that our statistical procedures were both replicable and generalizable to a replication data set. We 1) trained a portion of our data to identify predictive edges (i.e., 75% of discovery sample, the training data), 2) assessed the performance of the training set in a test portion of the training data (i.e., 25% of discovery sample, the test data), and 3) determined the generalizability of our results in an independent replication sample.

### A Framework for Future Investigations in Psychiatric Research

We also provide a statistical framework that can be used to assess the clinical utility of identification accuracy and to identify specific connections within and among brain networks of import. Viewing identification accuracy of rsfMRI scans as a classification problem is useful to other relevant questions in neuropsychiatric research. To improve early identification and detection of those at risk for psychiatric disorders, we need to answer questions such as, do people with similar connectivity profiles share common psychiatric features? Does reduced accuracy in functional connectome fingerprinting indicate risk for psychiatric disorders? These questions all fall within the realm of a classification problem, and the framework that we present here can be applied to relevant data to answer these questions.

There is work showing that distinct fingerprinting patterns map onto particular phenotypes or outcomes of interest. While, two studies report that fingerprinting accuracy is reduced in psychiatric samples [Kaufmann et al., 2017; Kaufmann et al., 2018], these were cross-sectional and relied upon comparison of already established clinical phenotypes (i.e., level of fingerprinting accuracy did not determine a *future* variable of interest). In the future, it will be important to use a sensitivity-specificity analytic framework to test our ability to identify sub-groups of individuals, such as adolescents who eventually go on to develop a psychiatric disorder or to predict treatment response in a group of patients.

### Limitations

As with any study, there are limitations present in our design. First, we split our sample by the median age and identified the two groups as “youths” and “adults” and this approach could obscure subtle differences in identification accuracy that occur across adolescent development. We chose this approach because identification accuracy increases with smaller sample sizes, and our sample size would be quite small if we split our sample into more age groups, as we have done in previous publications. Another approach could be to view age as a continuous variable; however, this imposes a strong linear assumption, when we know that development through adolescence is curvilinear in nature as stability is reached. In the future, examining identification accuracy in large samples of youth (i.e., the Adolescent Brain Cognitive Development Study, [Casey et al., 2018] in comparison to large samples of adults (e.g., Human Connectome Project, [Van Essen et al., 2013] may prove to be the most fruitful in terms of more fully understanding identification accuracy across development. Additionally, although we made all attempts remove noise in our processing steps (e.g., wavelet despiking and ICA-Aroma), we did not adjust for physiological noise (e.g., cardiac and respiratory cycles) in our analyses. Thus, we cannot rule out the possibility that this source affected our results. Physiological noise could be obscuring results in our data; it is even possible it could be generating part of the signal. In the future, it will be important to test fingerprinting accuracy after regressing out physiological influences or implementing analytic approaches to account for this noise in the data [Aslan et al., 2019].

### Conclusions and Future Directions

In this study, we showed that identification accuracy is high in both youths and adults at both short (same-day) and extended (1.5 years apart) periods of time. Importantly, our results suggest that networks properties may have an individualized foundational characterization that may be inherent to individuation with some room for flexibility in expression. In the future, we predict that combined use of group- and individual level data will become the sine qua non for identifying meaningful relationships between brain and behavior.

## References

Aslan S, Hocke L, Schwarz N, Frederick B (2019): Extraction of the cardiac waveform from simultaneous multislice fMRI data using slice sorted averaging and a deep learning reconstruction filter. Neuroimage 198:303–316.

Bathelt J, Johnson A, Zhang M, Astle DE (2019): The cingulum as a marker of individual differences in neurocognitive development. Sci Rep 9:2281.

Betzel RF, Bertolero MA, Gordon EM, Gratton C, Dosenbach NUF, Bassett DS (2019): The community structure of functional brain networks exhibits scale-specific patterns of inter- and intra-subject variability. Neuroimage 202:115990.

Calabro FJ, Murty VP, Jalbrzikowski M, Tervo-Clemmens B, Luna B (2019): Development of Hippocampal–Prefrontal Cortex Interactions through Adolescence. Cereb Cortex. https://academic.oup.com/cercor/advance-article-abstract/doi/10.1093/cercor/bhz186/5588470.

Casey BJ, Cannonier T, Conley MI, Cohen AO, Barch DM, Heitzeg MM, Soules ME, Teslovich T, Dellarco DV, Garavan H, Orr CA, Wager TD, Banich MT, Speer NK, Sutherland MT, Riedel MC, Dick AS, Bjork JM, Thomas KM, Chaarani B, Mejia MH, Hagler DJ Jr, Daniela Cornejo M, Sicat CS, Harms MP, Dosenbach NUF, Rosenberg M, Earl E, Bartsch H, Watts R, Polimeni JR, Kuperman JM, Fair DA, Dale AM, ABCD Imaging Acquisition Workgroup (2018): The Adolescent Brain Cognitive Development (ABCD) study: Imaging acquisition across 21 sites. Dev Cogn Neurosci 32:43–54.

Chawla NV, Bowyer KW, Hall LO, Kegelmeyer WP (2002): SMOTE: synthetic minority over-sampling technique. J Artif Intell Res 16:321–357.

DeLong ER, DeLong DM, Clarke-Pearson DL (1988): Comparing the areas under two or more correlated receiver operating characteristic curves: a nonparametric approach. Biometrics 44:837–845.

Demeter DV, Engelhardt LE, Mallett R, Gordon EM, Nugiel T, Harden KP, Tucker-Drob EM, Lewis-Peacock JA, Church JA (2019): Functional Connectivity Fingerprints at Rest are Similar Across Youths and Adults and Vary with Genetic Similarity. iScience:100801.

Dukart J, Bertolino A (2014): When structure affects function--the need for partial volume effect correction in functional and resting state magnetic resonance imaging studies. PLoS One 9:e114227.

Finn ES, Shen X, Scheinost D, Rosenberg MD, Huang J, Chun MM, Papademetris X, Constable RT (2015): Functional connectome fingerprinting: identifying individuals using patterns of brain connectivity. Nat Neurosci 18:1664–1671.

Florkowski CM (2008): Sensitivity, specificity, receiver-operating characteristic (ROC) curves and likelihood ratios: communicating the performance of diagnostic tests. Clin Biochem Rev 29 Suppl 1:S83–7.

Fonov VS, Evans AC, McKinstry RC, Almli CR (2009): Unbiased nonlinear average age-appropriate brain templates from birth to adulthood. NeuroImage.

Friedman J, Hastie T, Tibshirani R (2009): glmnet: Lasso and elastic-net regularized generalized linear models. R package version 1.

Galton F (1892): Finger Prints. Macmillan and Company.

Glasser MF, Coalson TS, Robinson EC, Hacker CD, Harwell J, Yacoub E, Ugurbil K, Andersson J, Beckmann CF, Jenkinson M, Smith SM, Van Essen DC (2016): A multi-modal parcellation of human cerebral cortex. Nature 536:171–178.

Gordon EM, Laumann TO, Adeyemo B, Huckins JF, Kelley WM, Petersen SE (2016): Generation and Evaluation of a Cortical Area Parcellation from Resting-State Correlations. Cereb Cortex 26:288–303.

Gordon EM, Laumann TO, Gilmore AW, Newbold DJ, Greene DJ, Berg JJ, Ortega M, Hoyt-Drazen C, Gratton C, Sun H, Hampton JM, Coalson RS, Nguyen AL, McDermott KB, Shimony JS, Snyder AZ, Schlaggar BL, Petersen SE, Nelson SM, Dosenbach NUF (2017): Precision Functional Mapping of Individual Human Brains. Neuron 95:791–807.e7.

Gratton C, Kraus BT, Greene DJ, Gordon EM, Laumann TO, Nelson SM, Dosenbach NUF, Petersen SE (2019): Defining Individual-Specific Functional Neuroanatomy for Precision Psychiatry. Biol Psychiatry. http://www.sciencedirect.com/science/article/pii/S0006322319318293.

Gratton C, Laumann TO, Nielsen AN, Greene DJ, Gordon EM, Gilmore AW, Nelson SM, Coalson RS, Snyder AZ, Schlaggar BL, Dosenbach NUF, Petersen SE (2018): Functional Brain Networks Are Dominated by Stable Group and Individual Factors, Not Cognitive or Daily Variation. Neuron 98:439–452.e5.

Greenwood JP, Maredia N, Younger JF, Brown JM, Nixon J, Everett CC, Bijsterveld P, Ridgway JP, Radjenovic A, Dickinson CJ, Ball SG, Plein S (2012): Cardiovascular magnetic resonance and single-photon emission computed tomography for diagnosis of coronary heart disease (CE-MARC): a prospective trial. Lancet 379:453–460.

Hafeman DM, Chase HW, Monk K, Bonar L, Hickey MB, McCaffrey A, Graur S, Manelis A, Ladouceur CD, Merranko J, Axelson DA, Goldstein BI, Goldstein TR, Birmaher B, Phillips ML (2019): Intrinsic functional connectivity correlates of person-level risk for bipolar disorder in offspring of affected parents. Neuropsychopharmacology 44:629–634.

Hallquist MN, Hwang K, Luna B (2013): The nuisance of nuisance regression: spectral misspecification in a common approach to resting-state fMRI preprocessing reintroduces noise and obscures functional connectivity. Neuroimage 82:208–225.

Horien C, Noble S, Finn ES, Shen X, Scheinost D, Constable RT (2018): Considering factors affecting the connectome-based identification process: Comment on Waller et al. Neuroimage 169:172–175.

Horien C, Shen X, Scheinost D, Constable RT (2019): The individual functional connectome is unique and stable over months to years. Neuroimage 189:676–687.

Hwang K, Hallquist MN, Luna B (2013): The development of hub architecture in the human functional brain network. Cereb Cortex 23:2380–2393.

Jalbrzikowski M, Larsen B, Hallquist MN, Foran W, Calabro F, Luna B (2017): Development of White Matter Microstructure and Intrinsic Functional Connectivity Between the Amygdala and Ventromedial Prefrontal Cortex: Associations With Anxiety and Depression. Biol Psychiatry 82:511–521.

Jalbrzikowski M, Liu F, Foran W, Roeder K, Devlin B, Luna B (2019): Resting-State Functional Network Organization Is Stable Across Adolescent Development for Typical and Psychosis Spectrum Youth. Schizophr Bull. http://dx.doi.org/10.1093/schbul/sbz053.

Jalbrzikowski M, Murty VP, Tervo-Clemmens B, Foran W, Luna B (2019, in press): Age-associated deviations of amygdala functional connectivity in youths with psychosis spectrum disorders: relevance to psychotic symptoms. Am J Psychiatry.

Kaufmann T, Alnæs D, Brandt CL, Bettella F, Djurovic S, Andreassen OA, Westlye LT (2018): Stability of the Brain Functional Connectome Fingerprint in Individuals With Schizophrenia. JAMA Psychiatry 75:749–751.

Kaufmann T, Alnæs D, Doan NT, Brandt CL, Andreassen OA, Westlye LT (2017): Delayed stabilization and individualization in connectome development are related to psychiatric disorders. Nat Neurosci 20:513–515.

Larsen B, Luna B (2018): Adolescence as a neurobiological critical period for the development of higher-order cognition. Neurosci Biobehav Rev 94:179–195.

Laumann TO, Gordon EM, Adeyemo B, Snyder AZ, Joo SJ, Chen M-Y, Gilmore AW, McDermott KB, Nelson SM, Dosenbach NUF, Schlaggar BL, Mumford JA, Poldrack RA, Petersen SE (2015a): Functional System and Areal Organization of a Highly Sampled Individual Human Brain. Neuron 87:657–670.

Laumann TO, Koyejo O, Gregory B, Hover A, Chen M-Y, Gorgolewski KJ, Luci J, Joo SJ, Boyd RL, Hunicke-Smith S, Simpson ZB, Caven T, Sochat V, Shine JM, Gordon E, Snyder AZ, Adeyemo B, Petersen SE, Glahn DC, McKay DR, Curran JE, Ring HHHGO, Carless MA, Blangero J, Dougherty R, Leemans A, Handwerker DA, Frick L, Marcotte EM, Mumford JA, Poldrack RA (2015b): Long-term neural and physiological phenotyping of a single human. Nat Commun 6:1–15.

Li R, Utevsky AV, Huettel SA, Braams BR, Peters S, Crone EA, van Duijvenvoorde ACK (2019): Developmental Maturation of the Precuneus as a Functional Core of the Default Mode Network. J Cogn Neurosci 31:1506–1519.

Luna B, Marek S, Larsen B, Tervo-Clemmens B, Chahal R (2015): An integrative model of the maturation of cognitive control. Annu Rev Neurosci 38:151–170.

Marek S, Hwang K, Foran W, Hallquist MN, Luna B (2015): The Contribution of Network Organization and Integration to the Development of Cognitive Control. PLoS Biol 13:e1002328.

Marek S, Tervo-Clemmens B, Nielsen AN, Wheelock MD, Miller RL, Laumann TO, Earl E, Foran WW, Cordova M, Doyle O, Perrone A, Miranda-Dominguez O, Feczko E, Sturgeon D, Graham A, Hermosillo R, Snider K, Galassi A, Nagel BJ, Ewing SWF, Eggebrecht AT, Garavan H, Dale AM, Greene DJ, Barch DM, Fair DA, Luna B, Dosenbach NUF (2019): Identifying reproducible individual differences in childhood functional brain networks: An ABCD study. Dev Cogn Neurosci 40:100706.

Miranda-Dominguez O, Feczko E, Grayson DS, Walum H, Nigg JT, Fair DA (2018): Heritability of the human connectome: A connectotyping study. Network Neuroscience 2:175–199.

Miranda-Dominguez O, Mills BD, Carpenter SD, Grant KA, Kroenke CD, Nigg JT, Fair DA (2014): Connectotyping: model based fingerprinting of the functional connectome. PLoS One 9:e111048.

Mueller S, Wang D, Fox MD, Yeo BTT, Sepulcre J, Sabuncu MR, Shafee R, Lu J, Liu H (2013): Individual variability in functional connectivity architecture of the human brain. Neuron 77:586–595.

Murty VP, Calabro F, Luna B (2016): The role of experience in adolescent cognitive development: Integration of executive, memory, and mesolimbic systems. Neurosci Biobehav Rev:1–32.

Patel AX, Bullmore ET (2015): A wavelet-based estimator of the degrees of freedom in denoised fMRI time series for probabilistic testing of functional connectivity and brain graphs. Neuroimage. http://eutils.ncbi.nlm.nih.gov/entrez/eutils/elink.fcgi?dbfrom=pubmed&id=25944610&retmode=ref&cmd=prlinks.

Paus T, Keshavan M, Giedd JN (2008): Why do many psychiatric disorders emerge during adolescence? Nat Rev Neurosci 9:947–957.

Power JD, Barnes KA, Snyder AZ, Schlaggar BL, Petersen SE (2012): Spurious but systematic correlations in functional connectivity MRI networks arise from subject motion. Neuroimage 59:2142–2154.

Power JD, Cohen AL, Nelson SM, Wig GS, Barnes KA, Church JA, Vogel AC, Laumann TO, Miezin FM, Schlaggar BL, Petersen SE (2011): Functional network organization of the human brain. Neuron 72:665–678.

Pruim RHR, Mennes M, Buitelaar JK, Beckmann CF (2015a): Evaluation of ICA-AROMA and alternative strategies for motion artifact removal in resting state fMRI. Neuroimage 112:278–287.

Pruim RHR, Mennes M, van Rooij D, Llera A, Buitelaar JK, Beckmann CF (2015b): ICA-AROMA: A robust ICA-based strategy for removing motion artifacts from fMRI data. Neuroimage 112:267–277.

Robin X, Turck N, Hainard A, Tiberti N, Lisacek F, Sanchez J-C, Müller M (2011): pROC: an open-source package for R and S+ to analyze and compare ROC curves. BMC Bioinformatics 12:77.

Satterthwaite TD, Wolf DH, Loughead J, Ruparel K, Elliott MA, Hakonarson H, Gur RC, Gur RE (2012): Impact of in-scanner head motion on multiple measures of functional connectivity: relevance for studies of neurodevelopment in youth. Neuroimage 60:623–632.

Shen X, Tokoglu F, Papademetris X, Constable RT (2013): Groupwise whole-brain parcellation from resting-state fMRI data for network node identification. Neuroimage 82:403–415.

Steinberg L (2005): Cognitive and affective development in adolescence. Trends Cogn Sci 9:69–74.

Stigler SM (1995): Galton and identification by fingerprints. Genetics 140:857–860.

Van Dijk KRA, Sabuncu MR, Buckner RL (2012): The influence of head motion on intrinsic functional connectivity MRI. Neuroimage 59:431–438.

Van Essen DC, Smith SM, Barch DM, Behrens TEJ, Yacoub E, Ugurbil K, WU-Minn HCP Consortium (2013): The WU-Minn Human Connectome Project: an overview. Neuroimage 80:62–79.

Waller L, Walter H, Kruschwitz JD, Reuter L, Müller S, Erk S, Veer IM (2017): Evaluating the replicability, specificity, and generalizability of connectome fingerprints. Neuroimage 158:371–377.

Xu T, Opitz A, Craddock RC, Wright MJ, Zuo X-N, Milham MP (2016): Assessing Variations in Areal Organization for the Intrinsic Brain: From Fingerprints to Reliability. Cereb Cortex. http://dx.doi.org/10.1093/cercor/bhw241.

Yeo BTT, Krienen FM, Sepulcre J, Sabuncu MR, Lashkari D, Hollinshead M, Roffman JL, Smoller JW, Zollei L, Polimeni JR, Fischl B, Liu H, Buckner RL (2011): The organization of the human cerebral cortex estimated by intrinsic functional connectivity. J Neurophysiol 106:1125–1165.

Yi C, Zeng Y (2018): sparseSVM: Solution Paths of Sparse High-Dimensional Support Vector Machine with Lasso or Elastic-Net Regularization. https://CRAN.R-project.org/package=sparseSVM.

Youngstrom EA (2014): A primer on receiver operating characteristic analysis and diagnostic efficiency statistics for pediatric psychology: we are ready to ROC. J Pediatr Psychol 39:204–221.

