## Supplemental Materials for "Functional connectome fingerprinting accuracy in youths and adults is similar when examined on the same day and 1.5 years apart"

**Supplementary Text and Tables**

**Discovery Sample: Detailed Exclusion Criteria**

In the discovery sample, the original sample size was 146 participants. One participant was removed due to poor registration and 5 more were removed for excessive motion (FD >0.5mm). The final sample for the discovery sample was N=140.

See Figure S1 for differing Ns at sessions and visits.

**Figure S1***.* Detailed exclusion criteria at sessions and visits in the discovery sample.

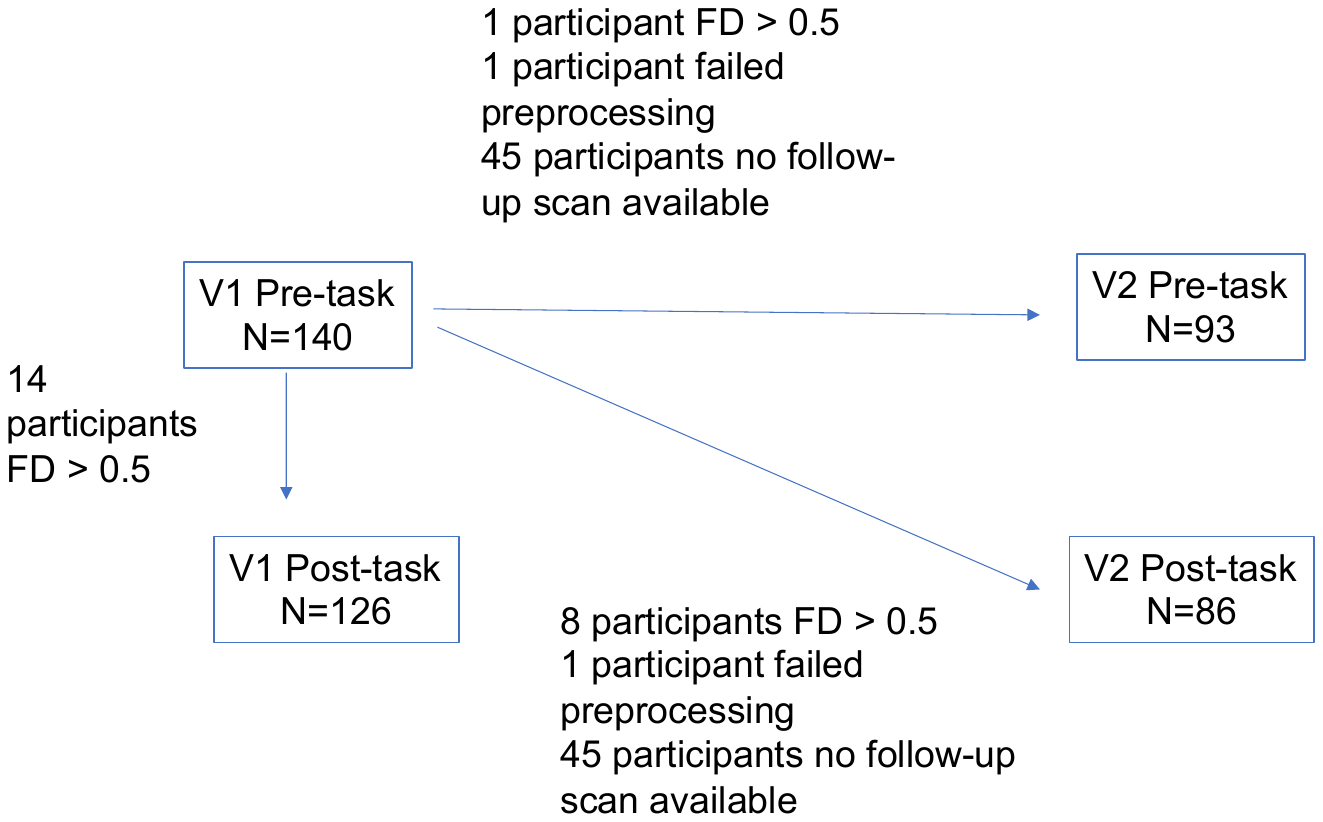

**Replication Sample: MR Data Acquisition**

Data were acquired using a Siemens 3 Tesla Tim Trio at the University of Pittsburgh Medical Center Magnetic Resonance Research Center using a 12-channel phase array head coil. We collected five minutes of resting-state data with eyes closed while awake. Functional images were acquired using an echo-planar sequence sensitive to BOLD contrast (T_2_*). rsfMRI parameters were: TR/TE=1500/29 ms, flip angle=70**°**, voxel size=3.125x3.125mm in-plane, 29 contiguous 4-mm axial slices, 200 TRs. A magnetization-prepared rapid gradient-echo sequence (MPRAGE) was acquired to measure brain structure and for alignment of the rsfMRI images. MPRAGE parameters were: TR/TE1570/3.4ms, flip angle=8°, TI=800 ms, voxel size:0.78125 x 0.78125 x 1 mm, 200 TRs.

**Figure S2**. Distributions of correlations between pairs from the discovery sample (red) and replication sample (blue) for the different methods: A) Finn, B) elastic net, C) SVM, and D) all edges.

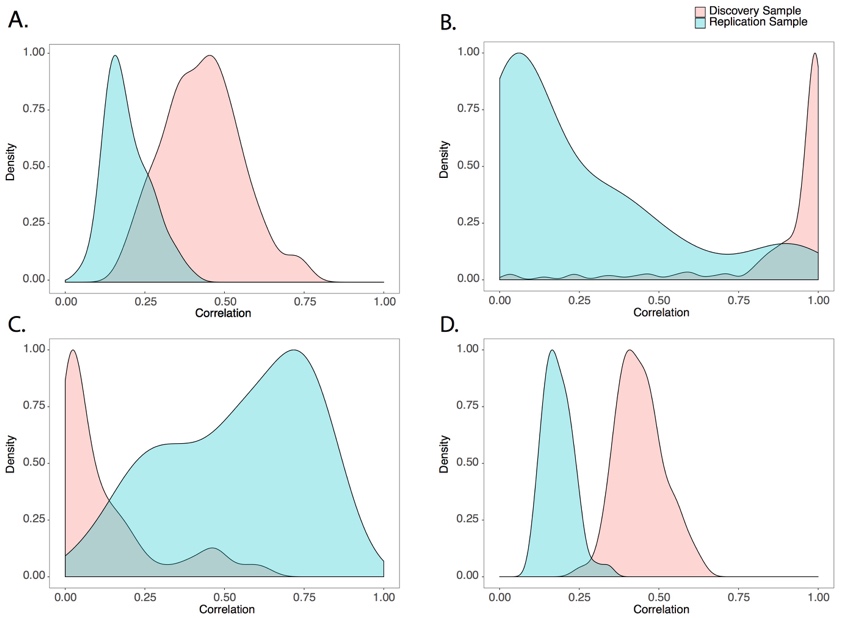

**Table S1**. rsfMRI sensory-motor and cognitive networks identified in the Gordon et al 2016 parcellation. Table details the number of nodes in each network, along with number of within and between network edges.

| Network | # of Nodes | # of Within-Network Edges | # of Between-Network Edges |
| --- | --- | --- | --- |
| Default | 41 | 820 | 11972 |
| Sensory-Motor Hand | 38 | 703 | 11210 |
| Sensory-Motor Mouth | 8 | 28 | 2600 |
| Visual | 39 | 741 | 11466 |
| Frontoparietal | 24 | 276 | 7416 |
| Auditory | 24 | 276 | 7416 |
| None | 47 | 1081 | 13442 |
| Cingulo-Parietal | 5 | 10 | 1640 |
| Retrosplenial-Temporal | 8 | 28 | 2600 |
| Cingulo-Opercular | 40 | 780 | 11720 |
| Ventral Attention | 23 | 253 | 7130 |
| Salience | 4 | 6 | 1316 |
| Dorsal Attention | 32 | 496 | 9632 |

**Table S2**. Demographic information of participants (age range: 18.1-30 years old) who had an additional MRI session later in the day.

| N | M/F | mean age | age range |
| --- | --- | --- | --- |
| 76 | 38/38 | 23.5 [3.7] | 18-31 |

**Table S3.** Demographic information for youth and adult groups in the training sample.

|  | **Pre-task** | | | | | **Post-task** | | | | |
| --- | --- | --- | --- | --- | --- | --- | --- | --- | --- | --- |
| **Youths** | N | M/F | Mean Age | Age Range | Mean FD | N | M/F | Mean Age | Age Range | Mean FD |
| **Visit 1** | 72 | 38/34 | 15.6 | 12.0-19.8 | 0.25 | 61 | 35/26 | 15.9 | 12.0-19.8 | 0.24 |
| **Visit 2** | 37 | 15/22 | 17.2 | 13.5-21.3 | 0.24 | 34 | 15/19 | 17.6 | 13.5-21.3 | 0.24 |
| **Adults** | N | M/F | Mean Age | Age Range | Mean FD | N | M/F | Mean Age | Age Range | Mean FD |
| **Visit 1** | 57 | 29/28 | 24.8 | 20.42-30.1 | 0.23 | 54 | 28/26 | 24.9 | 20.42-30.1 | 0.26 |
| **Visit 2** | 46 | 24/22 | 26.4 | 21.8-32.6 | 0.21 | 42 | 21/21 | 26.4 | 21.8-32.6 | 0.22 |

**Table S4.** Fingerprinting accuracy was similar in males and females separately.

| Sex | Visit/Session | AUC | Threshold | Sensitivity | Specificity |
| --- | --- | --- | --- | --- | --- |
| Female | Same-day V1 | 0.99 | 0.38 | 0.95 | 0.93 |
| Female | Same-day V2 | 0.97 | 0.40 | 0.90 | 0.91 |
| Male | Same-day V1 | 0.98 | 0.38 | 0.93 | 0.93 |
| Male | Same-day V2 | 0.95 | 0.41 | 0.87 | 0.93 |
| Female | 1 year pre-task | 0.90 | 0.35 | 0.84 | 0.80 |
| Female | 1 year post-task | 0.92 | 0.37 | 0.84 | 0.88 |
| Male | 1 year pre-task | 0.88 | 0.36 | 0.78 | 0.86 |
| Male | 1 year post-task | 0.90 | 0.37 | 0.84 | 0.85 |

**Table S5.** All three edge selection methods (Finn method, Elastic Net, and SVM) significantly improved identification accuracy in comparison to using all edges to identify an individual. These three methods performed similarly to one another.

| Comparison | D | *P* |
| --- | --- | --- |
| All edges vs. Finn method | -5.0 | 9.4e-07 |
| All edges vs. Elastic Net | -4.5 | 7.3e-06 |
| All edges vs. SVM | -4.8 | 1.4e-06 |
| Finn method vs. Elastic Net | 1.6 | 0.11 |
| Finn method vs. SVM | .6 | 0.43 |
| Elastic Net vs. SVM | -0.9 | 0.33 |

**Table S6.** In comparison to using all edges to identify an individual All three edge selection methods (Finn method, elastic net, and SVM) significantly improved identification accuracy in the test sample.

| Comparison | D | p-value |
| --- | --- | --- |
| All edges vs. Finn method | -4.8 | 1.3e-06 |
| All edges vs. Elastic Net | -4.6 | 4.5e-06 |
| All edges vs. SVM | -4.8 | 1.5e-06 |
| Finn method vs. Elastic Net | 0.39 | 0.70 |
| Finn method vs. SVM | 1.0 | 0.30 |
| Elastic Net vs. SVM | 0.07 | 0.94 |

**Table S7.** When predictive edges from the 1.5 year comparisons from the training sample were used to improve accuracy in the test sample, the finding remained similar to results obtained with all comparisons from the training sample (same day and 1.5 year comparisons). See Figure 4B for comparisons.

|  | AUC | Threshold | Sensitivity | Specificity |
| --- | --- | --- | --- | --- |
| Finn Method | 0.95 | 0.13 | 0.86 | 0.89 |
| Elastic Net | 0.94 | 0.02 | 0.79 | 0.93 |
| SVM | 0.94 | 0.26 | 0.86 | 0.85 |
| All Edges | 0.78 | 0.21 | 0.50 | 0.98 |

**Table S8. See Excel File,** Elastic Net weights obtained from training sample.

**Table S9. See Excel File,** SVM weights obtained from training sample.

**Table S10.** In all methods, the distribution of predictive/non-predictive edges for within-connectivity edges to the distribution of predictive/non-predictive edges to between-connectivity edges was significantly higher.

|  | X^2^ | p-value |
| --- | --- | --- |
| Finn method | 10.9 | 0.0009 |
| Elastic Net | 20.9 | 4.8E-06 |
| SVM | 72.5 | 1.6e-17 |

**Table S11.** See Excel file.

**Table S12.** Standardized residuals from chi-square test of over-representation of within-network connections when using the Finn method to detect predictive edges in same day and 1.5 year apart comparisons. Most predictive networks across comparisons are in bold. Least predictive networks across comparisons are italicized.

| Network | Same Day | | 1.5 years | |
| --- | --- | --- | --- | --- |
|  | V1 | V2 | Pre-Task | Post-Task |
| **Dorsal Attention** | **13.74** | **9.17** | **4.08** | **3.28** |
| **Frontoparietal** | **13.44** | **19.87** | **6.47** | **10.47** |
| **Default mode** | **4.25** | **8.31** | **4.16** | **11.16** |
| Salience | 2.09 | 2.47 | 2.7 | 2.59 |
| Cingulo-parietal | 1.76 | 2.17 | 2.42 | 2.3 |
| Ventral attention | 1.23 | -0.02 | 7.84 | 1.01 |
| Retrosplenial Temporal | 0.79 | 1.28 | 1.57 | 1.43 |
| Sensory-motor mouth | 0.79 | 1.28 | 1.57 | 1.43 |
| Visual | -1.42 | -6.34 | -3.49 | -4.41 |
| Auditory | -1.65 | -3.27 | -0.07 | -2.97 |
| Cingulo-opercular | -4.37 | -3.98 | -1.02 | -2.21 |
| *Sensory-motor hand* | *-6.38* | *-7.17* | *-5.87* | *-4.4* |
| *None* | *-11.21* | *-9.27* | *-6.7* | *-8.7* |

**Table S13**. With the Finn method, the distribution of predictive/non-predictive edges for within-connectivity edges to the distribution of predictive/non-predictive edges to between-connectivity edges was significantly higher in both youths and adults.

|  | X^2^ | p-value |
| --- | --- | --- |
| Finn method youths | 110.58 | 2.2e-16 |
| Finn method adults | 26.1 | 3.1e-07 |

**Table S14.** Standardized residuals from chi-square test of over-representation of within-network connections when using the Finn method to determine predictive edges in youth vs. adults separately. Most predictive networks across comparisons are in bold. Least predictive networks across comparisons are italicized.

|  | Youths | Adults |
| --- | --- | --- |
| **Frontoparietal** | **17.4** | **15.3** |
| **Dorsal Attention** | **12.5** | **6.4** |
| **Default** | **12.7** | **5.2** |
| Visual | -8.7 | 3.3 |
| Ventral Attention | 5.2 | -0.7 |
| Salience | -1.1 | -1.0 |
| Cingulo-opercular | -3.8 | -1.2 |
| Cingulo-parietal | -1.4 | -1.3 |
| Auditory | -6.2 | -1.6 |
| Retrosplenial Temporal | -2.2 | -2.0 |
| SMmouth | -2.2 | -2.0 |
| *SMhand* | *-6.9* | *-8.2* |
| *None* | *-11.6* | *-10.3* |

**Table S15.** See Excel file, Standardized residuals when testing over-representation of all between-network connections.

**Table S16.** See Excel file, Standardized residuals when testing over-representation of all network connections.

**Table S17.** Standardized residuals from chi-square test of over-representation of between-network connections when using the Finn method to detect predictive edges in same day and 1.5 year apart comparisons. Most predictive networks across comparisons are in bold. Least predictive networks across comparisons are italicized.

| Network 1 | Network 2 | Same Day V1 | Same Day V2 | Pre-Task 1.5 years | Post-Task 1.5 years |
| --- | --- | --- | --- | --- | --- |
| **Frontoparietal** | **Dorsal Attention** | **23.0** | **22.9** | **8.3** | **6.7** |
| **Frontoparietal** | **Default** | **21.5** | **20.9** | **18.3** | **26.1** |
| **Ventral Attention** | **Frontoparietal** | **13.9** | **13.5** | **11.3** | **9.1** |
| **Frontoparietal** | **Cingulo-opercular** | **13.9** | **14.0** | **10.2** | **6.9** |
| **Dorsal Attention** | **Default** | **12.2** | **14.3** | **5.9** | **8.8** |
| *None* | *Auditory* | *-5.9* | *-5.5* | *-4.7* | *-4.3* |
| *Sensory Motor Hand* | *Default* | *-5.9* | *-7.5* | *-6.3* | *-5.4* |
| *None* | *Cingulo-opercular* | *-7.6* | *-8.2* | *-4.5* | *-6.1* |
| *Visual* | *None* | *-8.4* | *-8.3* | *-5.7* | *-7.8* |
| *Sensory Motor Hand* | *None* | *-9.1* | *-9.3* | *-7.9* | *-8.4* |

**Table S18.** Standardized residuals from chi-square test of over-representation of between-network connections when using the Finn method to determine predictive edges in youth vs. adults separately. The five most predictive networks across comparisons are in bold. The five least predictive networks across comparisons are italicized.

| Network 1 | Network 2 | Youths | Adults |
| --- | --- | --- | --- |
| **Frontoparietal** | **Default** | **29.9** | **21.6** |
| **Ventral Attention** | **Cingulo-opercular** | **9.6** | **20.5** |
| **Frontoparietal** | **Dorsal Attention** | **16.4** | **17.5** |
| **Ventral Attention** | **Frontoparietal** | **11.9** | **16.9** |
| **Frontoparietal** | **Cingulo-opercular** | **10.4** | **15.3** |
| *None* | *Dorsal Attention* | *-5.3* | *-6.6* |
| *SMhand* | *Default* | *-8.1* | *-7.0* |
| *None* | *Cingulo-opercular* | *-5.0* | *-8.5* |
| *Visual* | *None* | *-5.7* | *-9.2* |
| *SMhand* | *None* | *-8.7* | *-9.4* |

**Table S19**. A) Within subject comparisons revealed that pre-task V1 framewise displacement was significantly higher than pre-task V2 framewise displacement. We also found that post-task V2 framewise displacement was significantly lower than post-task V1 framewise displacement. There were also no statistically significant differences in motion between B) youths and adults at each session.

| A. **Framewise displacement** | | |
| --- | --- | --- |
| Entire sample | T-value | p-value |
| pre-task V1 vs. pre-task V2 | 1.7 | 0.08 |
| post-task V1 vs. post-task V2 | 2.8 | 0.006 |
| pre-task V1 vs. post-task V1 | -2.5 | 0.01 |
| pre-task V2 vs. post-task V2 | -1.3 | 0.17 |
| Youth vs. Adults | T-value | p-value |
| pre-task V1 | 1 | 0.31 |
| post-task V1 | -1.8 | 0.09 |
| pre-task V2 | 2 | 0.05 |
| post-task V2 | 0.06 | 0.55 |
| B. **Mean head displacement** | | |
| Entire sample | T-value | p-value |
| pre-task V1 vs. pre-task V2 | 0.13 | 0.89 |
| post-task V1 vs. post-task V2 | 0.29 | 0.77 |
| pre-task V1 vs. post-task V1 | 0.10 | 0.92 |
| pre-task V2 vs. post-task V2 | -0.19 | 0.85 |
| Youth vs. Adults | T-value | p-value |
| pre-task V1 | -0.99 | 0.32 |
| post-task V1 | -1.19 | 0.24 |
| pre-task V2 | -0.30 | 0.76 |
| post-task V2 | -1.52 | 0.13 |
| C. **Maximum head displacement** | | |
| Entire sample | T-value | p-value |
| pre-task V1 vs. pre-task V2 | -0.08 | 0.93 |
| post-task V1 vs. post-task V2 | 2.3 | 0.02 |
| pre-task V1 vs. post-task V1 | -2.5 | 0.01 |
| pre-task V2 vs. post-task V2 | 0.37 | 0.71 |
| Youth vs. Adults | T-value | p-value |
| pre-task V1 | 1.47 | 0.14 |
| post-task V1 | 1.28 | 0.20 |
| pre-task V2 | 0.29 | 0.77 |
| post-task V2 | -0.48 | 0.64 |
| D. **# of Micromovements** | | |
| Entire sample | T-value | p-value |
| pre-task V1 vs. pre-task V2 | 1.08 | 0.28 |
| post-task V1 vs. post-task V2 | 2.85 | 0.005 |
| pre-task V1 vs. post-task V1 | -0.58 | 0.56 |
| pre-task V2 vs. post-task V2 | 1.39 | 0.16 |
| Youth vs. Adults | T-value | p-value |
| pre-task V1 | 1.03 | 0.30 |
| post-task V1 | 1.28 | 0.20 |
| pre-task V2 | 0.85 | 0.40 |
| post-task V2 | 0.56 | 0.58 |
